# A Proposed Unified Mitotic Chromosome Architecture

**DOI:** 10.1101/2021.10.14.464227

**Authors:** John Sedat, Angus McDonald, Herbert Kasler, Eric Verdin, Hu Cang, Muthuvel Arigovindan, Cornelis Murre, Michael Elbaum

## Abstract

A molecular architecture is proposed for an example mitotic chromosome, human Chromosome 10. This architecture is built on a previously described interphase chromosome structure based on Cryo-EM cellular tomography (1), thus unifying chromosome structure throughout the complete mitotic cycle. The basic organizational principle, for mitotic chromosomes, is specific coiling of the 11-nm nucleosome fiber into large scale approximately 200 nm structures (a Slinky (2, motif cited in 3) in interphase, and then further modification and subsequent additional coiling for the final structure. The final mitotic chromosome architecture accounts for the dimensional values as well as the well known cytological configurations. In addition, proof is experimentally provided, by digital PCR technology, that G1 T-cell nuclei are diploid, thus one DNA molecule per chromosome. Many nucleosome linker DNA sequences, the promotors and enhancers, are suggestive of optimal exposure on the surfaces of the large-scale coils.

**Significance Statement:** The significance of this proposed mitotic chromosome architecture is that a specific, sequenced chromosome, human Chromosome 10, can be built into a specific architecture that accounts for the dimensional values and cytological descriptions, a first time result. Since this molecular architecture is an extension of the interphase chromosome structure, a coiling of the 11-nm nucleosome fiber with further coiling, a unifying molecular structure motif is present throughout the entire mitotic cycle, interphase through mitosis.

## Introduction

Packing the DNA of a given chromosome, with multiple cm of DNA for an average human chromosome, into a mitotic chromosome (MC), several microns in length, requires a length compression (LC) of 10–20,000, a challenging problem. There are many models for MC architecture (reviewed in 4, 5), most emphasizing various sized loops (6), but most models do not fully account for the LC which is required to satisfy the packing density. In addition, most MC models do not account for the cytological modifications, seen in the familiar chromosome banding spreads, a requirement for an acceptable MC architecture.

This paper proposes a MC architecture that fully satisfies the LC, dimensional values, and cytological chromosome structure changes for an example chromosome, human Chromosome 10 (C10) (7). This chromosome, whose DNA is fully sequenced, has a DNA size of 46 mm that is packed into a MC (on average 5–6 microns in length), thus a LC of about 10,000 suggesting an important boundary condition for the architecture.

The proposed MC architecture is built on a previously described interphase chromosome architecture (1). In brief, a STEM Cryo-EM nucleus tomogram, preserving the interphase nuclear structures by the cryo procedures, and then suitably processed by deconvolution (8, but see 1, 9, 10) was made. Throughout the nucleus a 100–300 nm coiled nucleosome 11-nm fiber was documented. This interphase structure was termed a Slinky (see 2, motif cited in 3), and is described in detail in (1).

Table 1 summarizes the interphase chromosome structure. First, C10 DNA is depicted in Table 1A. Second, most DNA in the nucleus is organized into the familiar nucleosomes (NS) with extended linkers shown in Table 1B or with linkers compressed as hairpins to compact the NS 11-nm fiber. The NS can be rotated so the 11-nm face is oriented in the fiber direction as seen in Fig. 2A or Fig. 2C (as example), for space considerations. There are 669,000 NS in C10. Thirdly, the NS fiber is further coiled into 100–300 nm structures, defined as S; if we use 200 nm as diameter of S (the average diameter), then C10 compacted by S is ∼112 microns in length as shown in Table 1C. The S is coiled more tightly to form heterochromatin or pulled out variably for transcription.

**Table 1.**
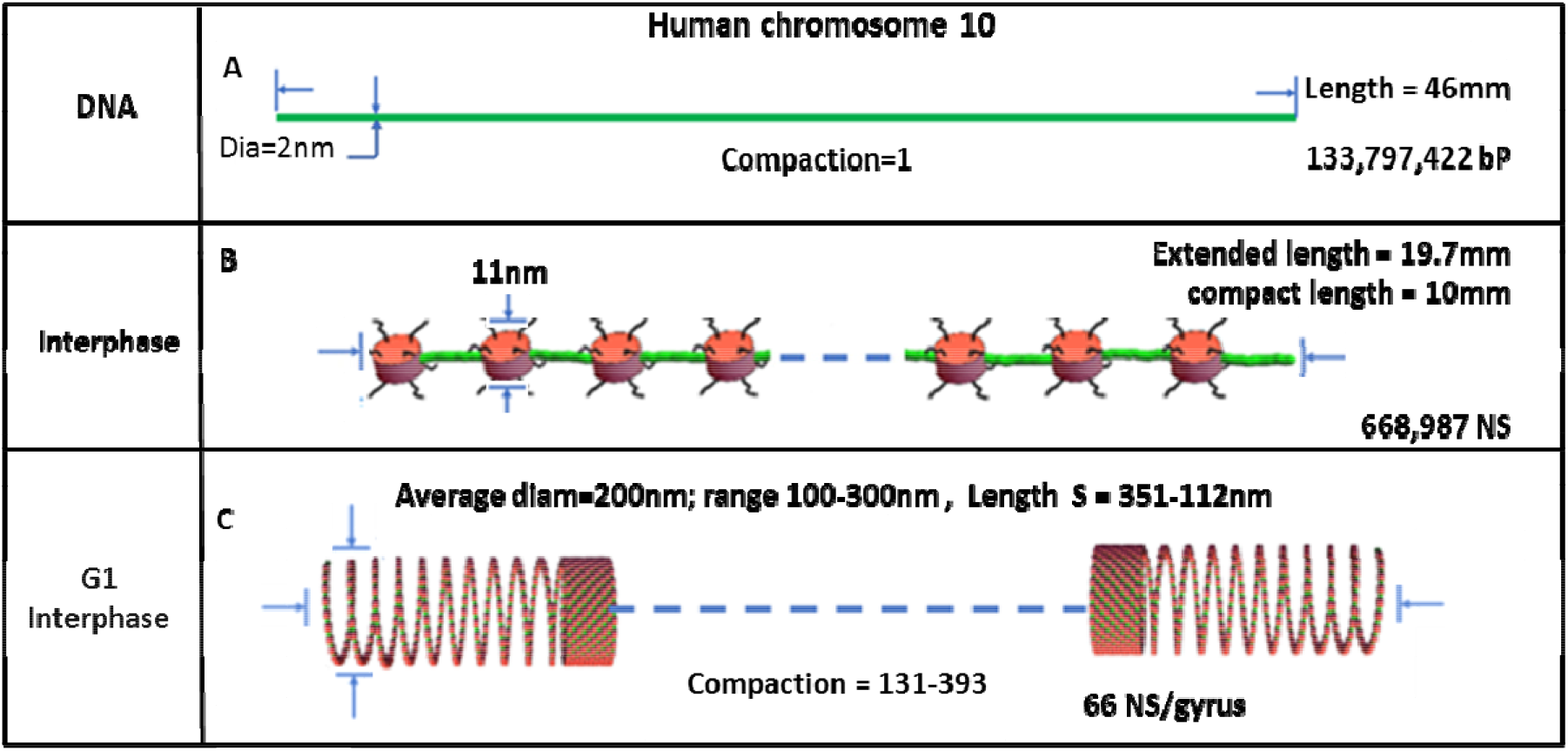
Table 1 documents C10 From DNA level to S NS Coil. Table 1A depicts C10 DNA, properly scaled. B shows the NS (scaled) with the green linker DNA extended while its possible the NS chain can be compressed with the linker DNA more compact as green hairpins. The histone tails are black wavy lines. C shows C10 NS coiling to form S with variable diameter, compaction and S length. The extended NS length is 669K NS (11nm [the NS size]) + 669K NS(18.4nm [linker DNA size]) while compact NS fiber is 669K NS [(11nm) +669K NS(2×2nm[DNA size for linker]) The number of S and S’ gyri can be calculated assuming 669K NS/ 66 NS/gyrus = 10,136 gyri. The compaction is S length / DNA length, where the S length is: 10mm[the compressed NS fiber length]/ S gyrus circumference [to give the number of S gyri] X 11nm([NS size; the gyrus size assuming maximal S packing].

This paper uses a package of software, described in the Methods, to sequentially coil multiple helices, for quantitative modeling, as a tool to explore mitotic chromosome architecture. This tool was also used in the interphase chromosome structure studies (1). The software models to an accurate (molecular) scale. Once mitotic chromosomes are built, they can be analyzed in any dimension or size scale.

The classical picture of the human genome (as a general example) is one of diploid organization, one DNA molecule per sister chromatid (11, 12). Nevertheless, there are still a few questions related to this generality, given the weak experimental data to support this conclusion (13). To clarify this boundary condition, a careful experimental protocol, using modern digital PCR technology with G1 T-Cells, was made, giving rise to a clear answer. The results show T-cells (in G1) are strictly diploid, hence one DNA molecule per single sister chromatid, the classical result.

## Results

### Direct Experimental Evidence That a Single Sister Mitotic Chromosome Has a Single DNA Molecule

We directly experimentally test the classical tenet that a single mitotic sister chromosome has only one copy of that chromosome’s DNA, or is uninemic (13). A stochiometric experiment was carried out. In brief, as described in the methods, multiple samples of T-cells in G1/G0, were cell-sorted. The T-cells were from a highly inbred mouse line, homozygous for all genetic loci, and as such had homologous chromosomes essentially with the same DNA sequence. The samples were spiked with a radioactive fragment of lambda DNA to control for losses, and the DNA extracted, making sure that the proteins did not mask DNA sequences. Carefully selected single copy genetic loci were used for Digital PCR analysis. Fig 1. Shows the results indicating that cells in G1/G0 have one copy of a DNA molecule per chromosome, a classical tenet in biology, and a boundary for the mitotic chromosome structure.

**Figure 1.**
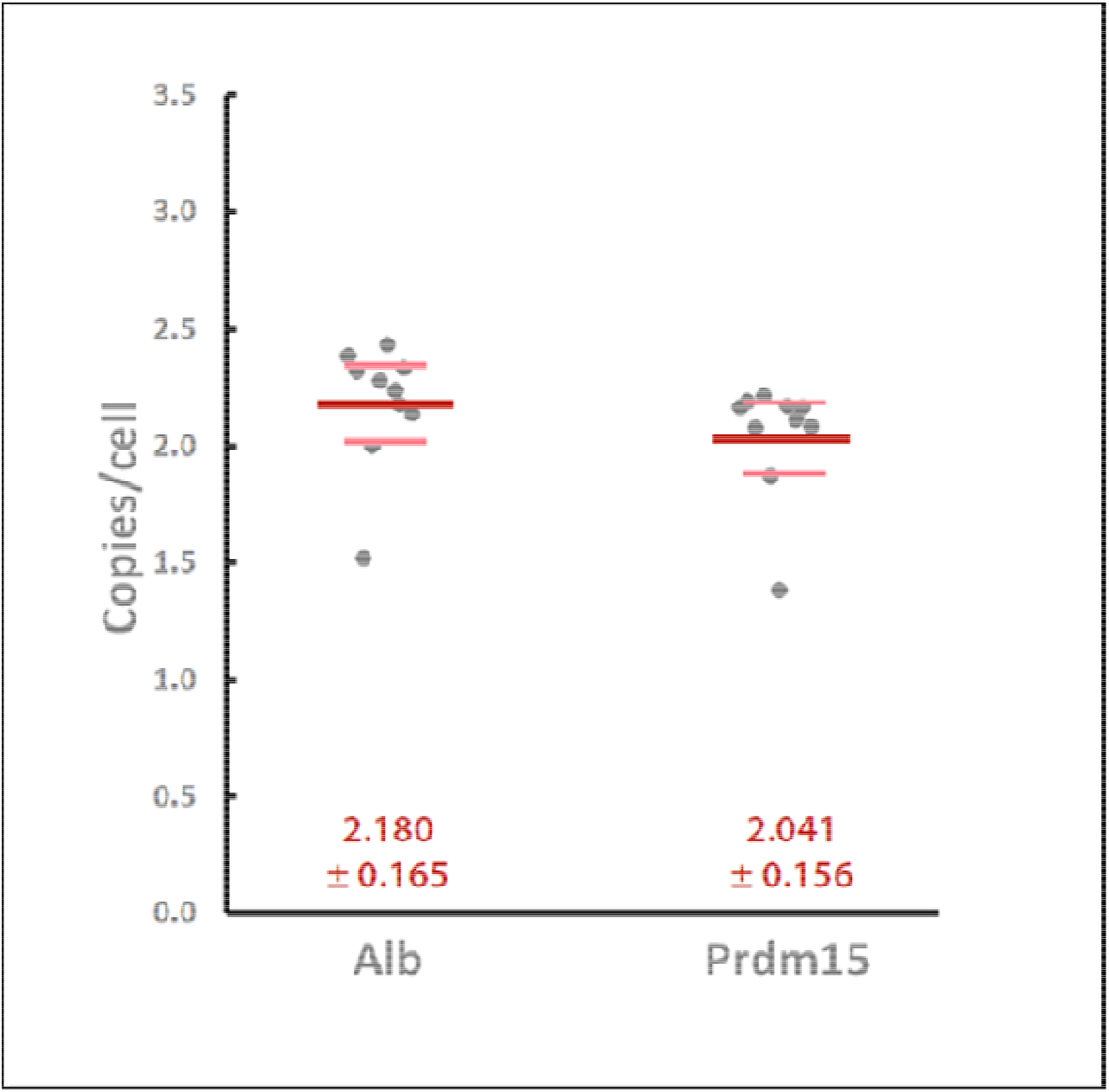
Copy number per cell (dark gray symbols) of two different genes that appear once per haploid genome, Albumin (left), and Prdm15 (right), as assessed by droplet digital PCR in pre-selection mouse CD4/CD8 double-positive thymocytes. Ten separate samples of 1 million pre-selection DP thymocytes (i.e. CD4^+^/CD8^+^/CD3^lo^/CD5^lo^) were flow sorted from two different C57bl/6 mice, and their genomic DNA prepared with radioisotope-based recovery tracking as described in Methods. A precisely known number of input cell equivalents were then amplified using primers and TaqMan probes specific for the indicated genes, and copies per cell calculated based on the counts of positive droplets. Bold red and lighter pink lines represent the means and 95% confidence intervals of the measurements respectively

**Figure 2.**
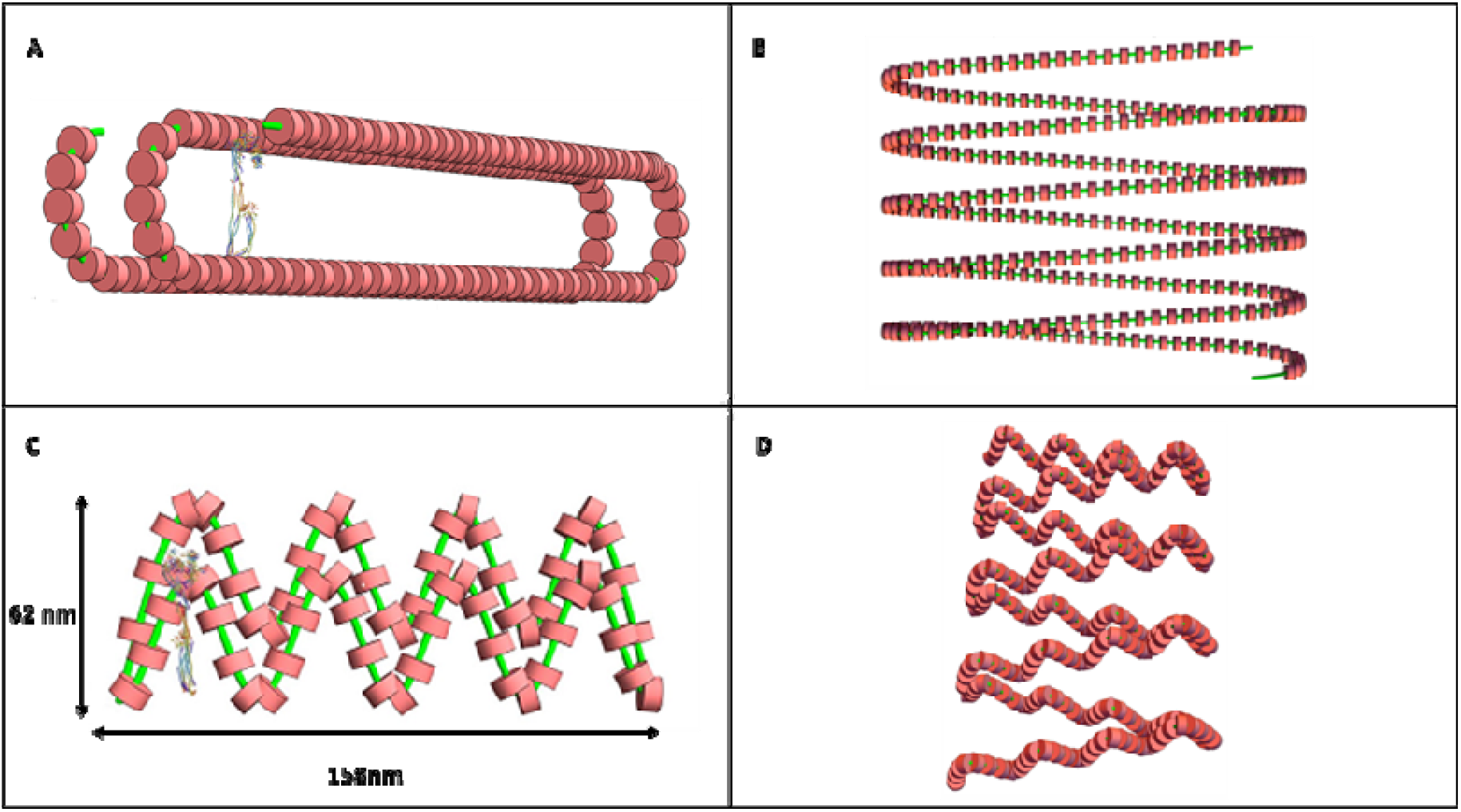
The S’ Coil is a Modified S Helix of the NS Fiber. Fig 2A depicts a single S’ RT gyrus, accurately to scale, with a Condensin I (23) for scale. Fig3B shows a RT S’ helical coiled region with coils more compressed. The minor axis of the RT is 50 nm with the major axis about 285 nm. Fig 3C shows a modified RT S’ further folded with short axis and long axis dimensioned. Fig 2D shows a modified S’ as a short coiled structure.

### S Modification Allows Further Interphase Chromosome Compaction: S’

Once an interphase chromosome S structure had been described (1), it was straight forward to model this structure, applying the coiling software (described in the Methods) with S to form a dimensioned C10 MC. The answer quickly showed that the compaction was off by an order of magnitude – only compacted 1,000 times – a common problem for MC architectures.

Study of the S cross-section shows that its hollow structure, on average 200 nm in diameter, was cylindrical and could be modified so its shape was indented, but still preserving the circumference, as shown in Fig. 2A. We noted that S could vary its diameter and NS packing (see Table 1B) as well as rotation of the NS (see Figs. 2 as examples), and we choose 66 NS/ gyrus as a reasonable packing for subsequent structure changes. We define this new S coil as S’ to reflect its modification. The cross-section of S’ can be variably compressed, reducing its short axis, but lengthening its long axis. The compressed S’ is an oval, shaped like a race track (RT); the ends have a circular geometry while the sides are straight lines, resulting in a short axis of about 50 nm and a long axis of approximately 290 nm. An S’ single gyrus is shown in Fig. 2A (& legend) with an extended S’ coil in Fig. 2B. The elongated cross-section could be (possibly) further compacted by helically twisting the structure shown in Fig. 2 (in essence a fat ribbon) so that it is a now very dense (reduced in cross-section size) rope – packed into a roughly circular, dense structure. This aspect of the structure was not further modeled into the MC. The RT, as shown in Fig. 2A, was incorporated into the modeling software, as described in the Methods, for the S’ architecture.

An ellipse was also considered for the compression of S into S’. A problem with the S’ ellipse are the tight turns at the top of the ellipse; the NS are differentially compressed. Some of the compression can be taken up by the flexible extendable NS linker DNA, but a more interesting modification is to reduce the compression by rotation of the NS by 90 degrees so the NS is now a 5-nm disk wedge (instead of a 11-nm disk) in the plane of the NS fiber, better accommodating the tight turn. The modeling software did not have this flexibility. The ellipse also had a reduced interior volume, restricting protein occupation. The ellipse was abandoned for the S’ structure.

The S’ RT was further considered for additional structures, primarily further coiling of S’. However, further coiling showed that the centers of such coiling were very dense, so modifications of the S’ RT (now MRT) were necessary. Fig. 2C shows that additional folds could be indented into the MRT – in essence creating loops –, further compressing the long axis of the RT, and the S’ coil of such structure is shown in Fig. 2D. As shown below, such a structure now allowed a mitotic chromosome to be built.

### The S’ Structure Can Be Further Coiled to Form the Mitotic Chromosome

The S’ structure depicted in Fig. 2D can be further helically coiled, about the long axis of the new MRT, to additionally compact the structure into a mitotic chromosome. This new additional coiling is shown in Fig. 3 as a half turn of the coil (Fig. 3A), and as two and a half turns (gryri) of the helical twists (Fig. 3B). These are properly dimensioned, and the NS of the previous winds are visible. We define this additional coiling S’’.

**Figure 3.**
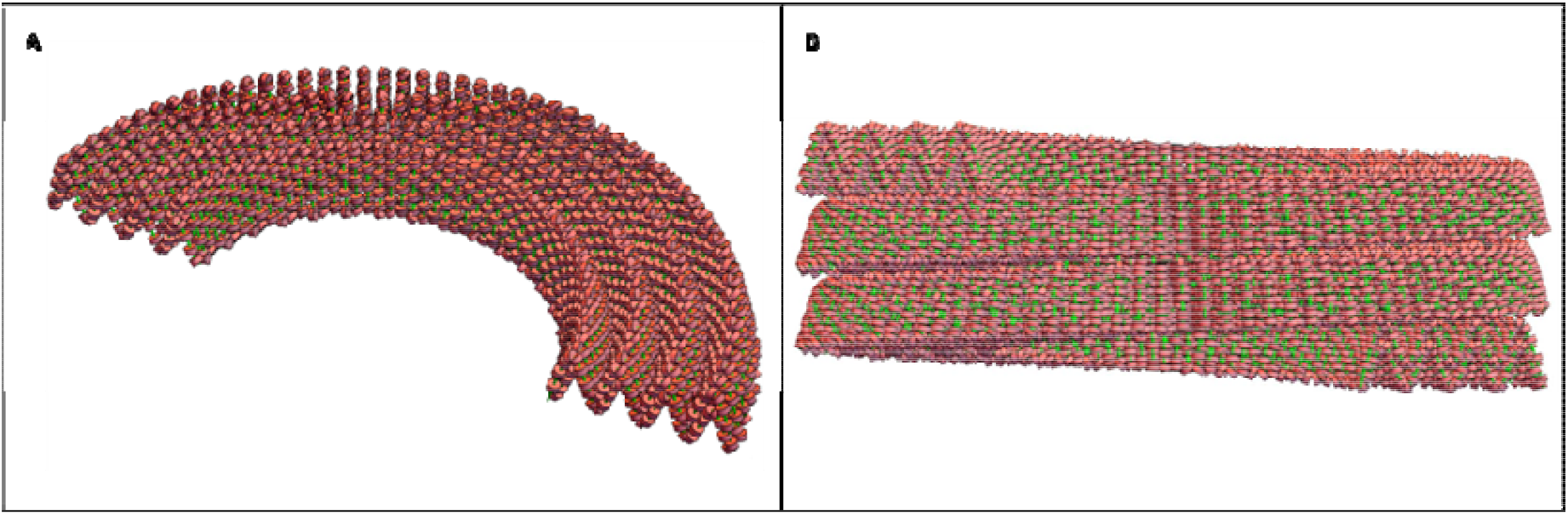
S’ Coils can be Further Helically Coiled to Form a S’’ Helix, The Final Coil of the MC. One half helical turn of the S’’ using the modified S’, drawn to scale, is shown (Fig 3A). The width of the S’ coil (the minor axis of the MRT) is ∼ 60 nm, and the S’’ coil is the width of the MC, approximately 0.6 microns. Fig 3B shows two and a half helical turns of S’’. NS of the S’ coils are visible.

It is now possible to build with the S’’ structure the C10 mitotic chromosome as shown in Fig. 4A. In order to accommodate the 669,000 NS in C10, there are ∼97 (96.5) full helical turns of S” to be made, resulting in a chromosome length of about 6 microns by ∼0.6 (0.59) microns diameter. This length/diameter and the S’’ NS packing satisfies the required LC, while the length/diameter matches the observed approximate cytological chromosome spread lengths/widths (see ref. 7). Fig. 4B shows the cross-section of the MC with its hollow center. We note the very densely packed NS. Thus, a credible MC architecture results.

**Figure 4.**
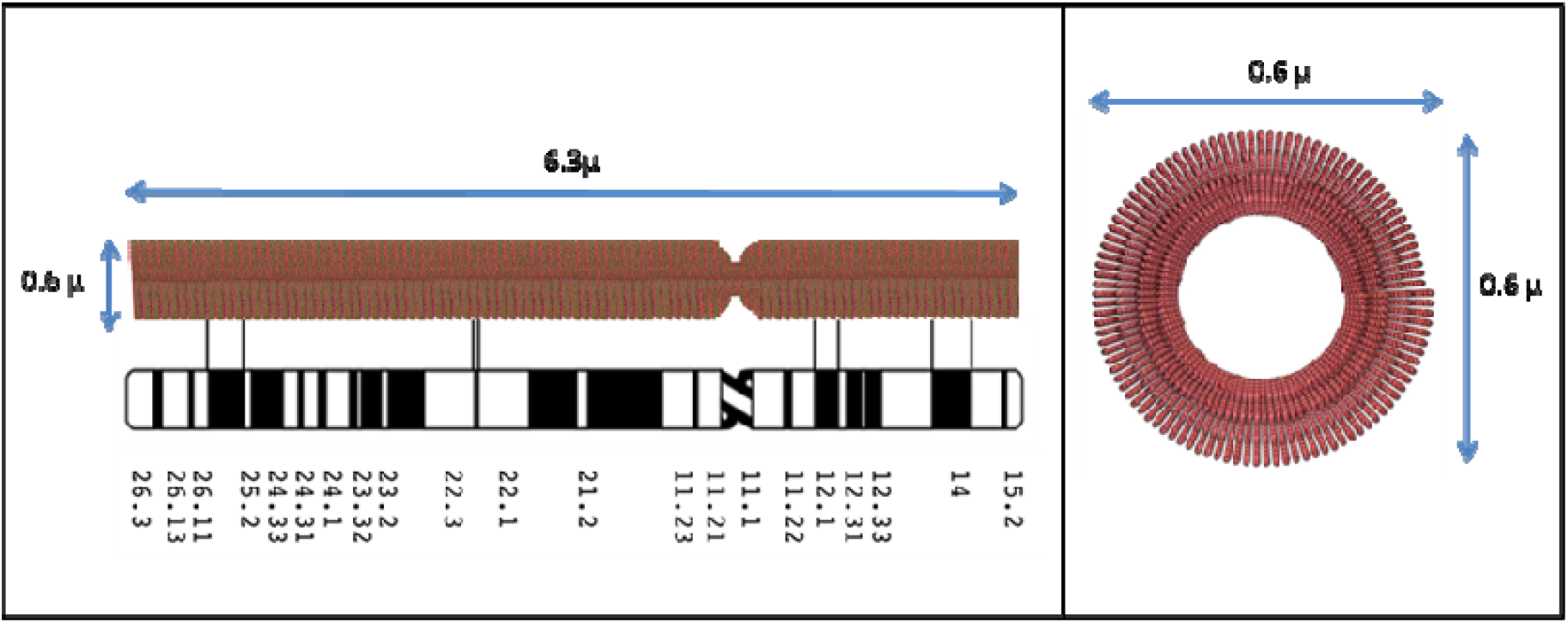
C10 Mitotic Chromosome structure. The left panel shows 97 S’’ helix winds/coils that build the C10 MC for a length of about 6 microns and a diameter of about 0.6 microns. The centromere constriction separates the two unequal arms (scaled) of C10, a metacentric MC. The lower panel depicts the banding cytology map for C10 (7), as accurately aligned with the structure as possible. The right panel is a cross-section of C10 MC, properly dimensioned.

## Discussion

There are several points to bring out in a discussion of the proposed MC architecture: First, there is a link of the architecture to the well-known MC cytology spreads (see 7) and chromosome manipulation procedures to compact some regions while decondensing other regions that stain differentially in a reproducible manner. Genetic locations, transcription events as well as deletions, while function at the DNA level, are localized on the MC bands (7). The banding patterns – broad as well as thin bands – for C10 are shown in Fig. 4. What is seen is that there is a straight boundary line clear across the MC width, delineating the stained band from the decondensed MC regions. We conjecture that the bands reflect the underlying coils of S’’; for example, the thin bands are one turn (gyrus) (about the right size), while a broad band is an integral number of gyri of S’’. Sister Chomatid Exchange (SCE), a recombination phenomenon (14), is another example of cytology and architecture linkup. SCE procedures differentially stain parental chromatids so that mitotic recombination exchanges are readily visualized (15). The exchanges show dark bands reciprocally moved to light sister chromatids, but the boundary of the dark/light exchange bands cuts straight across the MC width, like the above cytological banding patterns (see 14). The sharp dark/light lines, we conjecture, are the boundaries of S’’ gyri. All these features of our MC model reflect an essential feature of MC architecture, its colinearity with its DNA.

Secondly, the MC architecture possibly positions transcriptional events optimally for the next G1. The faces of the S’’ gyri have densely packed NS linker DNA sequences, the promotors and enhancers sequences, as shown in Fig. 5. The MC cytology banding patterns described above suggest that many related genetic pathways are located on specific bands (7). We propose that the faces of the S’’ gyri would expose the NS regions in a more easily searched fashion (in early G1) to optimize the transcription process sequence.

**Figure 5.**
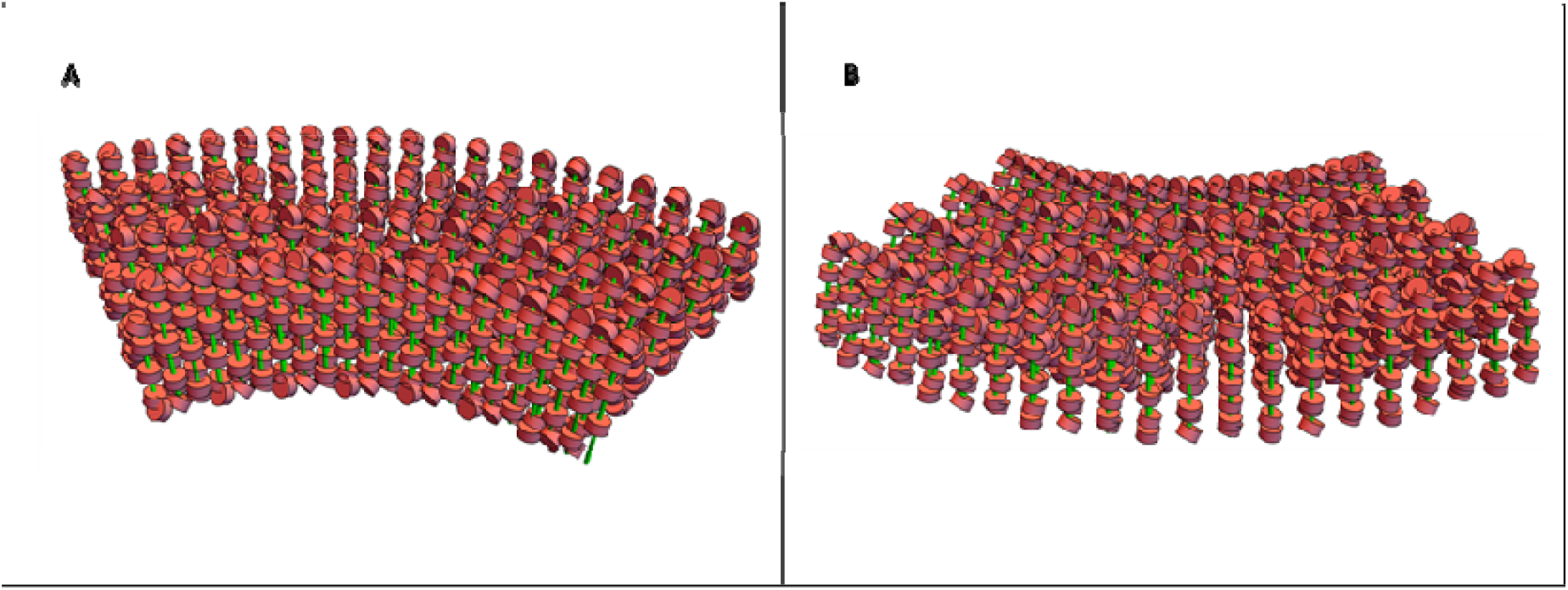
Faces of the S’’ winds highlight optimally exposed NS linker DNA. Fig 5A shows the enlarged inside surface of the S’’ half coil wind. See Fig 3 for orientation. The enlarged back face of the S’’ wind is shown in Fig 5B.

Thirdly, while we build the MC architecture in a specific, almost rigid crystalline fashion, the reality is that there is likely a great deal of architectural flexibility. First, the NS, shown in Table 1B, have linkers extended or compact which would extend (or compact) the NS fiber. Second, different numbers of NS could be coiled in S, changing the S diameter, a fact that is observed in the experimental data (1). Then, S’ could be modified by changes in the major/minor axis of the RT, the extreme of which is shown in Fig.2C&2D. This would change the packing density. Third, S’’ could be coiled more tightly or loosely to change the MC structure. Finally, pulling out S’ and S’’ in various places (changing the helical pitch even for short distances), also changes the MC structure. All coiling modifications could be working in concert in various combinations. The many chromosomal proteins (16) are likely distributed differentially along the length of the MC, and they might have a differential packing effect on the MC (see 7,16), further modifying the structure. We make no statement as to handedness; essentially all these levels could have a specific left- or right-hand coil, and that hand could in principle flip, though the transition region would be locally disorganized.

We are stunned by the density of the NS packing that is required to satisfy the boundary conditions. It is almost crystalline in density! Every packing feature, to increase the packing density, has to be used to get the structure to fit the MC, LC, and sizes. There is little room for slop.

We are at the upper end of the dimensions of MC C10, and reduction of the dimensions might be possible. There is flexibility in the structures for accommodation in the tight packing density. In the tight turns, Fig. 2C and inside the MC hole (Fig. 3 and Fig. 5A), for example, the substantial flexible linker DNA on each side of the NS will allow optimal movement. Flipping the NS by 90 degrees (now 5.5 nm in size), at the ends of the short axis of the MRT, will allow for tighter S’’ coiling and reduction of MC diameter and length. The pitch of S’’, or the number of S’ gyri/ S’’ gyrus will change the packing density/MC size also.

It is formally possible that some (or many) NS are removed from the S or S’ structures in the compacting stages of the MC so significant bare DNA can pack more densely. No evidence exists for this possibility. We are careful to state that other MC architecture schemes are possible, but not explored.

Fourthly, while we are careful to not say how such coiling is molecularly specified for this MC architecture, such coiling is used in many places in cell biology, such as in the structure of Myosins (11). The abundant MC proteins (16) are likely to make possible such coiling.

The major chromosomal proteins, the Condensins I & II (17–22), can be brought into the architecture, and the MRT S’ structure lends itself for this protein interaction. The latest Condensin I structure from the PDB (19) has a major axis of 34–36 nm which comes close to matching the new RT short axis of about 50 nm as seen in Fig. 3A and 3B (drawn to scale). Condensins I & II, similar in structure, are shaped to interact with, bind, and pull together the sides of the short axis of the RT as discussed in (17–22). A stick figure model of Condensin taken from (20) can also dock in the RT (Fig. 3A) for comparison. Cohesin, a very similarly shaped molecule (23, 24) could also interact with the RT. A publication describes a careful study of the stoichiometry of Condensin I & II throughout the cell cycle (21). Indeed, a large hole in the center of the S’’ and the MC results from this architecture matching the data of (21). There are other places for key proteins to bind and aid in the structure; Histone H1 (25, 26) could bind two NS pairs, above and below -in trans- the multiple folds in the modified S’ (Fig. 2C), which could consolidate/strengthen this structure. In summary, the RT MC structure suggests that a protein scaffold would be an essential component of this architecture.

Fifth, it is formally possible that several (two to four) of the S’’ gyri associate, in phase, through protein associations, to form larger structures possibly of the order of 0.2–0.3 microns (see Fig. 2). These larger structures are seen in some mitotic chromosome preparations (27), including many in the Sedat laboratory (unpublished) over the years.

Sixth, the very dense packing that is required to compress the NS into a MC suggests that biochemical perturbations such as fixation or chromosome isolation buffers (see 5) are a major problem. This very high density of packing will facilitate crosslinking, bringing structures artificially together while pulling other structures apart with a distorted final structure. The solution will likely be structure-preserving Cryo-EM tomography.

Seventh, the proposed MC architecture, with its dense closely packed helices, emphasizes that rigorous structure biology interpretation tools will be required to adequately solve, with proof, the structure. One can easily get lost among the multiple coils, especially with reduced resolution or not adequate Z resolution, leading to a false final structure. For an example, it is difficult, even with stereo viewing, to convince oneself that the final S’’ coiling of the MC is not a two stranded structure, though we know by building the structure that this cannot be true. The final MC structure awaits study by Cryo-EM tomography.

### Major Conclusion

Finally, in summary, we emphasize that a monolithic helical coiling architecture is used throughout the entire mitotic cell cycle from interphase through mitosis. Table 2 outlines a summary of the architecture as a function of the cell cycle. Table 2A shows the base level, the C10 DNA starting point. Table 2B shows the interphase G1/S cell cycle next level, the extended NSs. In Table 2C are shown the interphase G1/S NS, organized into S with a 100–300nm diameter, either pulled out for transcription or compacted for heterochromatin. In Table 2D, the prophase cell cycle level is shown with S indented to form S’, thus starting to further compress the NS for additional packing. Finally, Table 2E shows the mitotic cell cycle level with the MC coiled into S’’ helix. This architecture satisfies the packing boundary conditions, for example LC and dimensions, we outlined in the introduction. In addition, we can conclude, with some certainty, that one haploid DNA molecule forms one sister MC. Thus, there is a unity, as outlined in this proposal, for chromosomal structure—with one monolithic coiling architecture--throughout the entire cell cycle.

**Table 2.**
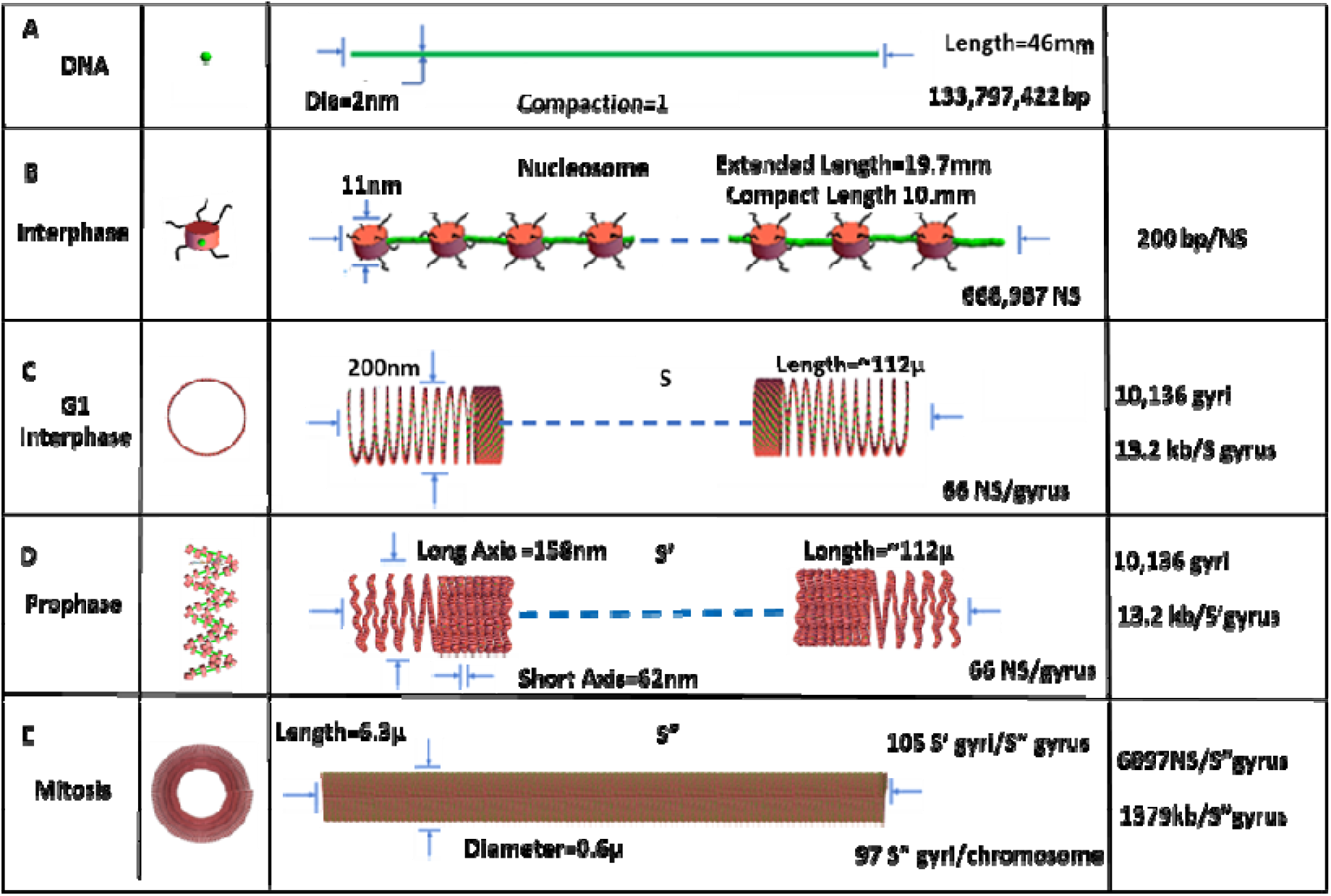
A Summary of the Unified Architecture Changes, using the Helical Multiple Coiling Motif Taking Place Throughout the Cell Cycle, Interphase through Mitosis a complete cell cycle, for C10. Table 2A is the DNA level, and Table2B shows the NS 11nm organization with extended linker DNA (green) Table 2C depicts the S level with the coiled NS tightly packed with 66 NS/gyrus, while Table 2D indents /folds the S structure to a MRT S’. Table 2E is the final level of organization, a mitotic chromosome based on S’’ coiling. The structures are drawn to scale. See Table 1. legend. The 6897NS/S’’ gyrus is given by 668,987 NS/97 S’’gyri

## Materials and Methods

### G1 copy number determination

Suspensions of total thymocytes at 2×10^7^/ml in PBS/2% FCS were prepared from two different male 8 week-old C57bl/6 mice by crushing the organs, straining through a 40 micron nylon mesh, and Ack lysis of erythrocytes. The cells were then stained with Antibodies (eBioscience) to mouse CD4 (clone GK1.4, PE), CD8 (clone 53-6.7, PE-Cy7), CD3 (clone 2C11, APC-AF780), and CD5 (clone 53-7.3, APC) at 1 µg/ml each for 20 minutes on ice. After washing and resuspension in PBS/2% FBS, single pre-selection thymocytes (CD4+/CD8+/CD5^lo^/CD3^lo^) were sorted into pre-weighed (0.1mg precision) 15 ml conical tubes at exactly 1 million cells/tube, using a BD FACSAria II cell sorter. The tubes were weighed again after sorting, and then a ^32^P-labeled bacteriophage λ-based recovery tracking probe was added to the cells, with the exact number of counts added (≈ 11,000–12,000 CPM, ≈ 50 µl) determined by the weight of the added probe solution.

The recovery probe was prepared as follows: A standard 100 µl PCR reaction was prepared containing 10 ng of bacteriophage λ DNA (NEB), 5 units of Taq polymerase (NEB), and 5 pmoles each of the following primers: TACGAACGCCATCGACTTACGCGTGCGC (5’ end) and GCCATTGCTCAGGTCGAAGAGATGCAGG (3’ end), which amplify a 6,086 bp segment of the lambda genome. The crude PCR product was gel purified from 1% agarose by spin column (Qiagen), and 200 ng of the purified amplicon were end-labeled using 10 µCi of γ-^32^P ATP (Amersham) and T4 polynucleotide kinase (NEB). The labeling reaction was then desalted on a G-25 spin column and further precipitated with 70% ethanol, after addition of 20 µg of glycogen carrier (Roche), to remove any unincorporated label. The probe was resuspended in TE-buffer at 249 cpm/mg solution at the time of addition, and the subsequent recovery determinations were adjusted for the decay of a reference sample.

After probe addition, the sorted cells (≈ 2 ml volume) were adjusted to 1% SDS, 20 mM Na^+^-EDTA, and 40 units/ml proteinase K (Roche), and incubated overnight at 55°C. The digested lysate was then incubated at 80°C for 45 minutes to inactivate the proteinase K, then precipitated with 70% ethanol, air dried, and resuspended in ≈ 200 µl of TE-buffer. The tubes were weighed again to determine the mass of the resuspended DNA solution, then a ≈ 20 µl sample from this was weighed and counted, based on which the % recovery of each sample after digestion and precipitation was determined.

Based on this determination, 2,928–4,231 cell equivalents per droplet generation reaction (0.81 µl/sample, in triplicate) were assayed using the Bio-Rad QX100 digital PCR system, for the copy number of two different genes, Alb and Prdm15, each of which occurs only once in the mouse genome. Sufficient droplets were acquired for each reaction to give a 95% confidence interval of less than ±5% for template concentrations according to the Poisson distribution.

Primers for the ddPCR analysis were as follows: Alb fwd: GTTACCAAGTGCTGTAGTGGAT Alb rev: GTGCAGATATCAGAGTGGAAGG, Alb probe: ACTGTCAGAGCAGAGAAGCATGGC, amplicon length = 128; Prdm15 fwd: ATGGATGTGGTCCCTGAGTA, Prdm15 rev: CCTGTCGGAGCAACATGAA, Prdm15 probe: CGCAGGTGTACTTCTTGTCACCGT, amplicon length = 113.

### Computer modeling Software

The computer modeling of the MC, at all levels, utilized a software package written by a Turkish Engineering group (28) that allows sequential helical coiling of defined sized structures. This package is run under, and requires, Mathematica 12.3 (29) under Linux in large work stations. Mathematica 12.3 was run under Windows 7, Intel Core i5 CPU 3.20 GHz processor with 4 cores and displayed on a Samsung C27F591monitor and NVIDIA Quadro K1200 video card and 8 GB of memory. Mathematica was also run on a CentOS 7.3 Linux system running on a Intel Core i7 CPU 2.80 GHz processor with 16 GB of memory. The display of the Linux generated output was visualized on the Windows 7 computer. The software was modified to take into account the RT S’ configuration modifications. The various scripts for the software are supplied by the authors upon request.

### Computer Display Software

Once structures are built they are displayed, in various dimensions, with the generalized display and quantitation software package written over the years by the Agard/Sedat groups (30, unpublished extensions by Eric Brandlund). This software, Priism, and its extensive Help files are available through the Agard and Sedat UCSF e-mails. The display scripts are also available from the authors.

## Acknowledgments

We are grateful for the advice and input from professors Marc Shuman, Markus Noll, Lloyd Smith, Robert Stroud, Elizabeth Blackburn, and David DeRosier. CM acknowledges grant ROIA 100880.

